# The effects of selfing, limited seed dispersal and environmental persistence on the evolution of size-dependent sex allocation

**DOI:** 10.64898/2026.11.23.768536

**Authors:** Samuel Michel, Charles Mullon, Thomas Lesaffre

**Affiliations:** Department of Ecology and Evolution, University of Lausanne, 1015 Lausanne, Switzerland; Department of Biology, University of Fribourg, 1700 Fribourg, Switzerland

**Keywords:** Sexual systems, Phenotypic plasticity, Environmental heterogeneity, Hermaphrodites

## Abstract

Sex allocation, the division of resources between male and female function, often varies as a function of size in cosexual plants. This variation is thought to evolve because fitness returns gained from allocating to each function are influenced by plant size, and many mechanisms have been proposed for why it should be so. Here, we explore how competing selfing and limited seed dispersal jointly shape the evolution of size-dependent sex allocation in an annual plant population where size is environmentally determined. First assuming that the quality of habitat patches changes every generation, we show that competing selfing favours increased femaleness with size, whereas limited seed dispersal selects in the opposite direction. Which of the two mechanisms prevails depends on the selfing and seed dispersal rates, as well as on the severity inbreeding depression. Then, we let patch-quality differences persist over several generations and show that such persistence generates an additional selective pressure on sex allocation when seed dispersal is limited. Environmental persistence favours larger plants to allocate more resource to female function, as producing more seeds increases the probability that their own offspring settle on their high-quality patch in the next generation.

## 1 Introduction

Most plants are cosexual, meaning that individuals produce both ovules and pollen (Renner, 2014). However, relative investment in female versus male function, hereafter sex allocation, often varies between individuals within populations. In particular, sex allocation has been shown to vary with plant size in many species, with larger plants often investing relatively more resources to their female function (Wright and Barrett, 1999; Paquin and Aarssen, 2004; Cao and Kudo, 2008; Zhao et al., 2008; Chen and Pannell, 2025a; Cai et al., 2026, see De Jong and Klinkhamer, 1989 and Klinkhamer et al., 1997 for a review), although the opposite situation where larger plants are more male has also been documented in several cases (e.g., Paquin and Aarssen, 2004; Delesalle and Mazer, 2009; Nakahara et al., 2018). Hereafter, we refer to these two patterns of allocation, where larger individuals are more female or more male, as “feminisation” and “masculinisation”, respectively (Fig. 1 for illustration). Such size-dependent sex allocation is thought to be favoured because the fitness returns gained from allocating resources to male and female function vary with plant size (Klinkhamer et al., 1997; Zhang and Jiang, 2002; Cadet et al., 2004, Chap. 9 in De Jong and Klinkhamer, 2005).

**Figure 1:**
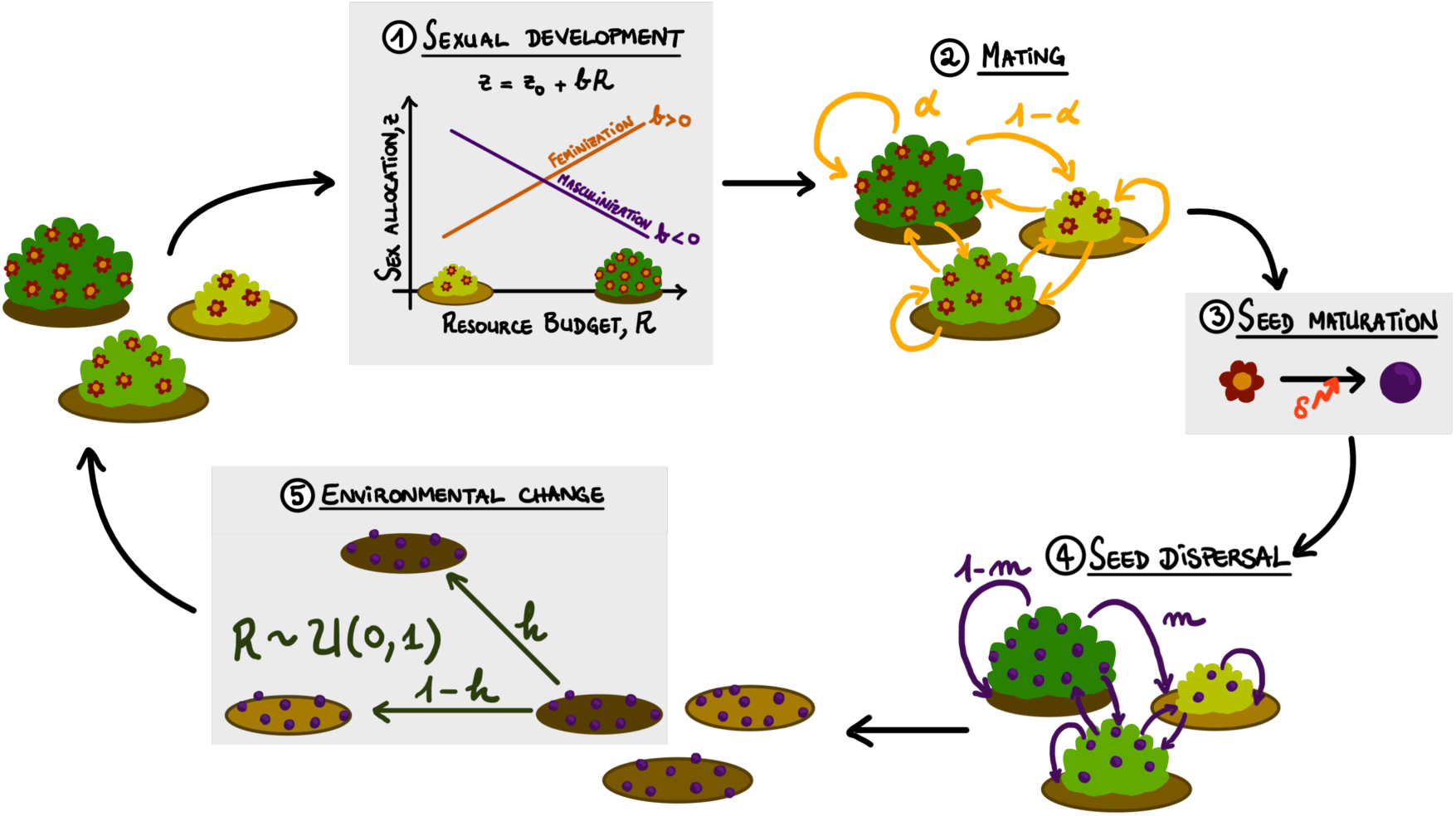
Life cycle followed by the population. Individuals develop on patches of varying quality. They first mature sexually and produce pollen and ovules according to their resource budget and sex allocation strategy. feminisation occurs when larger individuals allocate more to their female function (*b >* 0, orange) and masculinisation occurs when they allocate more to their male function (*b <* 0, purple). Individuals then exchange pollen following a mass-action model, retaining a fraction *α* of their pollen for self-fertilisation. This results in the production of diploid zygotes. self-fertilised zygotes suffer from inbreeding depression, surviving to maturity with probability 1 - *δ*, whereas all outcrossed zygotes become viable. Viable zygotes divide to produce two haploid seeds each.

Plant size can affect the fitness returns of ovule and pollen production in different ways that have been classified as either direct or budget effects (Klinkhamer et al., 1997). Direct effects come about when plant size itself affects expected fitness returns per ovule or pollen grain (Cadet et al., 2004), for instance if larger plants disperse pollen more efficiently. Meanwhile budget effects, on which we focus here, arise because size is often correlated with the resource budget available for reproduction: larger plants produce more gametes in absolute terms, and this in itself can influence fitness returns per gamete.

One important source of budget effects is the limited dispersal of pollen and seed. Limited pollen dispersal leads related pollen grains to compete for access to the same ovules (“local mate/sperm competition” Hamilton, 1967; Schärer, 2009; Fromhage and Kokko, 2010; Mullon et al., 2026), and limited seed dispersal causes related seedlings to compete for the same resources (“local resource competition”, Clark, 1978; Fromhage and Kokko, 2010; Mullon et al., 2026). Such kin competition among a plant’s own pollen grains or seeds is expected to intensify as its pollen or ovule production increases, which generates diminishing fitness returns to increased allocation to male and female function. Larger, more fecund individuals are therefore expected to shift their sex allocation towards the more efficiently dispersed sexual function, for which fitness returns diminish less rapidly (Sakai and Sakai, 2003).

Apart from kin competition under limited dispersal, self-fertilisation (hereafter selfing) – a common mating strategy in flowering plants whereby individuals fertilise their own ovules (Goodwillie et al., 2005) – is known to strongly affect relative fitness gains through male and female function (e.g., Charlesworth and Charlesworth, 1981; De Jong et al., 1999; De Jong and Geritz, 2001; Lesaffre et al., 2026a), and is also expected to generate a budget effect when the selfing rate depends on individual pollen production (as found in several species, e.g., Harder and Barrett, 1995; Vrieling et al., 1999; Chen and Pannell, 2024, 2025b). De Jong et al. (1999) investigated this idea through mathematical modelling and showed that large plants should invest relatively more resource to their female function when the selfing rate increases with pollen production. They suggested that this pattern is due to the greater difficulty for large plants to avoid geitonogamous selfing (i.e., selfing due to pollen transfers between the flowers of the same plant), which would in turn select for female-biased sex allocations. De Jong et al. (1999) assumed that the selfing rate depends only on the individual’s own pollen production. This corresponds to a form of prior selfing (Lloyd, 1979), in which self-fertilisation occurs before any opportunity for outcrossing. How other modes of selfing, and notably competing selfing in which self- and cross-pollen compete for ovule fertilisation, affect the evolution of size-dependent sex allocation remains unknown (Lloyd, 1979; Holsinger, 1991; Goodwillie et al., 2005). Importantly, De Jong et al. (1999) did not consider the effect of inbreeding depression, which is known to crucially impact the evolution of plant mating and sexual systems and might therefore alter the size dependence that evolves (Lande and Schemske, 1985; Goodwillie et al., 2005; Charlesworth and Charlesworth, 1981; Freeman et al., 1997; Charlesworth, 1999; Pannell and Jordan, 2022).

In this paper, we use mathematical modelling to study the evolution of size-dependent sex allocation under competing selfing and in the presence of inbreeding depression. We consider an annual plant population evolving in a patch-structured habitat where each patch hosts a single adult plant, the size of which is determined by the quality of the patch. As a baseline, we assume that patch quality changes randomly each generation and that all seeds are dispersed away from their natal patch. Under these assumptions, we show that competing selfing favours feminisation provided that inbreeding depression remains low (so that selfing is beneficial). Under high inbreeding depression, meanwhile, selection for inbreeding avoidance favours the emergence of separate sexes (De Jong and Geritz, 2001).

We then examine how two additional factors might interact with the budget effect created by competing selfing in shaping the evolution of size-dependent sex allocation. First, we consider the effect of limited seed dispersal. Our analysis reveals that the masculinising effect of limited seed dispersal often outweighs the feminising effect of selfing in the presence of inbreeding depression. This leads masculinisation to evolve over a broader range of parameters, especially when inbreeding depression approaches a half. When inbreeding depression exceeds a half, limited dispersal counteracts selection for inbreeding avoidance and opposes the emergence of separate sexes, leading to strong masculinisation. Where sexual polymorphism does nonetheless emerge, either separate sexes or gynodioecy (the coexistence of females and hermaphrodites) may evolve depending on parameters.

The second factor that we consider is the possibility that differences in patch quality persist over several generations. Such environmental persistence may be a common feature of plant populations. For example, some patches of habitat may provide better access to sunlight, water or nutrients for extended periods of time because of slowly changing features of the landscape, such as topography or undersoil composition. When seed dispersal is limited, environmental persistence increases the probability that seedlings experience the same growing conditions as their mother. This can favour feminisation, because larger plant growing in good conditions can increase the probability that their offspring inherit their high-quality patch by allocating more to female function (Chapter 9 in De Jong and Klinkhamer, 2005, Guillon et al., 2006). However, previous models describing this effect assume infinitely large habitat patches, so that there is no kin competition (Guillon et al., 2006). As a result, it remains unclear how the feminising effect of environmental persistence compares with the masculinising effect of local resource competition. We find that local resource competition predominates at low seed dispersal rates, leading to masculinisation. Conversely, environmental persistence dominates a higher dispersal rates, leading to feminisation. Taken together, our results highlight that the evolution of size-dependent sex allocation is influenced by several interacting factors, and that the direction and strength in which they act may depend on the fine-scale structure of the habitat the population evolves in.

## 2 The model

We consider a population of annual hermaphrodites evolving in an environment composed of *N* patches that can host one adult plant each. The patches differ in quality, offering a uniformly distributed resource budget *R* ∈ [0, 1]. Each generation, the resource budget that each patch offers remains the same with probability *h* or takes a new random value with probability 1 − *h*. The parameter *h* thus captures the persistence of environmental heterogeneity across generations. We assume that individuals growing on high-quality patches reach a higher size than those growing on lower quality ones. Hereafter, we therefore refer to plant growing in better conditions as larger for ease of presentation. Each generation, individuals pass through the following life-cycle events (Fig. 1).

### (i) Sexual development

Individuals first mature sexually and allocate a proportion *z* ∈ [0, 1] of their resource budget to the production of ovules and the rest to the production of pollen. We assume that the sex allocation *z* may depend on the resource budget *R*, so that the sex allocation of individual *i* with a resource budget *R_i_* is given by

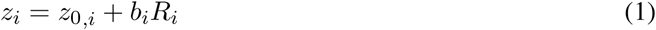

where *b_i_*∈ R and *z*_0_*_,i_* ∈ R are genetically encoded traits. Component *z*_0_*_,i_* determines the baseline sex allocation of individual *i*, whereas the slope component *b_i_* determines how sex allocation changes with the individual’s resource budget. Thus, individuals display size-dependent sex allocation whenever *b* ≠ 0: a positive slope *b >* 0 indicates that plants show feminisation, and a negative slope *b <* 0 indicates that they show masculinisation (because *b* and *z*_0_ can take any real value, we clip sex allocation *z* to ensure that it always remains in the [0, 1] interval, see Appendix A for details). We assume that phenotypic plasticity can be costly, so that the amount of ovules and pollen produced by individual *i* are given by

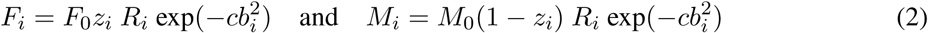

Individuals then disperse a fraction *m* of their seeds to other patches, and the remainder 1 − *m* falls on their natal patch. Each patch remains of the same quality with probability *h*. Otherwise, a new resource budget is sampled in a uniform distribution, *R* ∼ U[0, 1]. Finally, seeds compete on each patch to form the next generation.

where *F*_0_ and *M*_0_ are constants that give the number of ovules and pollen grains produced per unit of resource invested in each function, and *c >* 0 measures the cost of phenotypic plasticity, which could for example stem from the need to sustain the machinery required to sense the environment or maintain developmental flexibility (DeWitt et al., 1998). We assume throughout that *F*_0_ is large (specifically that *F*_0_ ≫ *N*) so that the population always remains at carrying capacity, and that *M*_0_ is larger (*M*_0_ ≫ *F*_0_), so that there is always enough pollen for all ovules to be fertilised (no pollen limitation).

### (ii) Mating

After gamete production, individuals keep a fraction *α* ∈ [0, 1] of the pollen they produced for self-fertilisation and export the rest (1 − *α*) to fertilise ovules on other plants. The exported pollen is evenly distributed across the *N* − 1 other individuals in the population. The selfing rate *s_i_* of individual *i* is determined by the proportion of self-pollen out of the total amount of pollen that it received on its stigma

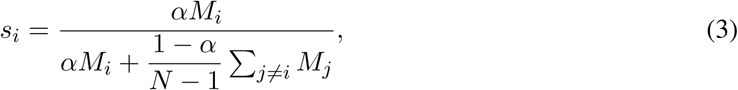

which corresponds to a mass-action model of competing selfing (Holsinger, 1991). The selfing rate of individual *i* thus depends on its resource budget and sex allocation as well as those of other individuals in the population.

### (iii) Seed maturation

Fertilisation results in the production of diploid zygotes. Outcrossed zygotes become viable with probability 1 and then immediately divide to give rise to two haploid seeds each. self-fertilised zygotes, meanwhile, suffer from inbreeding depression, which reduces their chances of surviving and dividing into mature haploid seeds by a proportion *δ* ∈ [0, 1]. Plants are therefore assumed to be haploid with a reduced diploid phase in our model. This assumption lets us investigate the evolution of size-dependent sex allocation in the absence of genetic constraints, so that any combination of baseline *z*_0_ and slope *b* components may evolve with in principle no impact on long-term evolutionary outcomes (e.g., Geritz and Kisdi, 2000; Lesaffre et al., 2025; Lehmann and Mullon, 2026; Lesaffre et al., 2026a).

### (iv) Seed dispersal

Following seed maturation, individuals disperse a fraction *m* ∈ [0, 1] of the seeds they produced towards other patches and the rest of the seeds (1 − *m*) remain on their natal patch. Dispersed seeds may fall on any of the *N* − 1 other patches with equal probabilities.

### (v) Seed germination, recruitment and environmental change

All adults die after seed dispersal, and the seeds present on each patch compete locally for recruitment. The quality of each patch may then change. Specifically, the amount of resource provided by patch *i* in the next generation either remains unchanged (with probability *h*), or is drawn in a uniform distribution *R* ∼ U[0, 1] (with probability 1 − *h*).

We assume the baseline and slope components of sex allocation *z*_0_ and *b* to be controlled by two fully linked quantitative trait loci. Alleles at these loci can encode any value of *b* and *z*_0_ (i.e., the loci follow a “continuum-of-alleles” model; Kimura, 1965). Each generation, alleles have a small probability *µ* to mutate, creating a new allele that deviates from the ancestral allele by a small amount sampled in a centred normal distribution with standard deviation *σ_µ_*.

We analysed the model using individual-based simulations implemented in C++. A detailed description of the simulation program can be found in Appendix A, and the code and output data are publicly available at 10.5281/zenodo.24459591.

## 3 Results

### 3.1 Competing selfing favours either feminisation or the emergence of separate sexes

We first investigated a baseline scenario where seed dispersal is complete (*m* = 1) and patch quality does not persist across generations (*h* = 0), so that only selfing and inbreeding depression influence the evolution of sex allocation (Section 3.1). In this case, evolutionary outcomes depend on the severity of inbreeding depression.

*Low inbreeding depression (δ <* 1*/*2). When *δ <* 1*/*2, plasticity evolves such that the population displays feminisation (*b >* 0, Fig. 2A). This is because selfing is net-beneficial here, as the transmission advantage that comes from self-fertilising one’s seeds outweighs the reduction in their viability that it causes, so that selfed seeds transmit more gene copies than outcrossed ones on average (Fisher, 1941; Lande and Schemske, 1985). It is thus advantageous for a plant to increase the number of self-fertilised seeds it produces. This number is limited both by the plant’s selfing rate and its ovule production, which vary in opposite directions with plant sex allocation. Individuals must therefore strike a balance between these two factors, and this balance differs depending on the resource budget: larger plants can achieve the same selfing rate with a smaller fraction of their resource budget spent on pollen, and can therefore allocate more resources to ovule production. To confirm this, we ran additional simulations where the selfing rate was fixed and equal for all individuals. As expected, we found that size-dependent sex allocation did not evolve in this case (i.e., the slope component remained close to *b* = 0 throughout), and only the baseline component *z*_0_ evolved to a female-biased value (Supp. Fig. S1) as expected in the absence of plasticity (Charlesworth and Charlesworth, 1981)).

**Figure 2:**
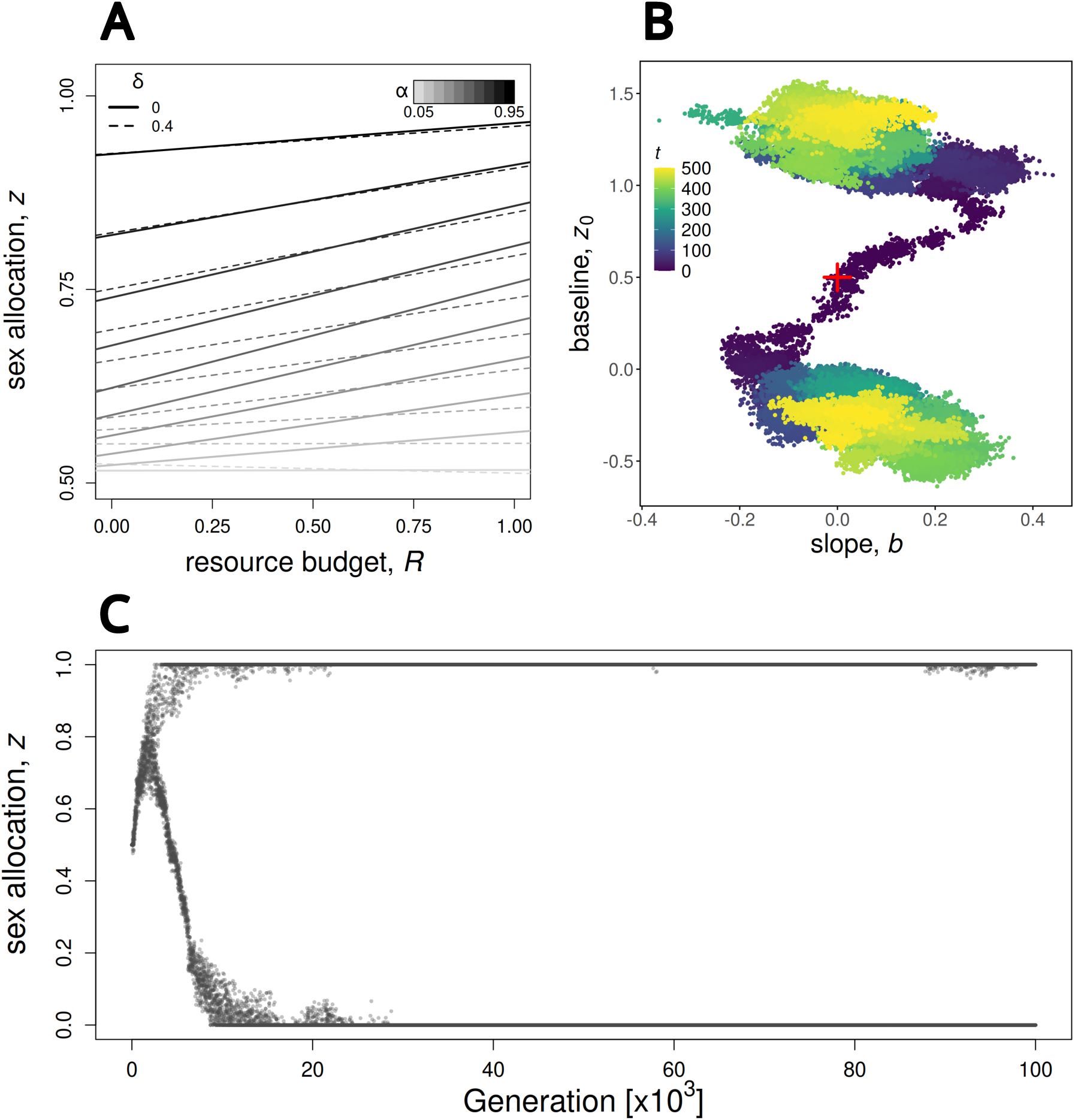
Effect of selfing and inbreeding depression on the evolution of size-dependent sex allocation. (A) Evolved size-dependent sex allocations under weak inbreeding depression (*δ <* 1*/*2). Each line represents the evolved size-dependent sex allocation as a function of resource budget for a different combination of *α* and *δ*. Darker colours indicate higher selfing rates (higher values of *α*). Continuous and dashed lines show evolved sex allocations in the absence of inbreeding depression (*δ* = 0) and under moderate inbreeding depression (*δ* = 0.4), respectively. For each parameter combination, we ran three replicates for 2 × 10^6^ generations each. The reaction norms shown here were obtained by averaging slope and baseline components from a hundred randomly sampled individuals every hundred generation over the last 5 × 10^5^ generations, across replicates. (B) Evolutionary trajectory of sex allocation components in a sample simulation under high inbreeding depression and selfing (*δ* = 0.6 and *α* = 0.75). Each point represents an allele, defined by its slope (x-axis) and baseline (y-axis) components. The colour gradient indicates time forward in the simulation from darker to lighter. The plot only shows the first 5 × 10^5^ generations for readability. (C) Distribution of phenotypes through time in the same simulation as in panel B for the first 10^5^ generations. Each point is an individual randomly sampled from the population. Parameters used in all simulations: *N* = 5, 000, *µ* = 0.005, *σ_µ_* = 0.01, *m* = 1, *h* = 0, *c* = 0.01.

Our simulations also revealed an effect of the severity of inbreeding depression on the evolution of size-dependent sex allocation: increasing inbreeding depression *δ* (while still maintaining it below *δ* = 1*/*2) reduces the magnitude of size dependence (i.e., reduces the slope *b*; Figure 2A). This is because increasing inbreeding depression reduces the viability of selfed seeds and thus the benefits of producing them. In turn, this reduces the benefits of size dependence relative to its cost (as measured by the cost of plasticity *c* in eq. 2), leading to weaker size dependence at evolutionary equilibrium.

*High inbreeding depression (δ >* 1*/*2). When *δ >* 1*/*2, the population undergoes a transition from combined to separate sexes: selection for inbreeding avoidance leads female- and male-biased alleles to emerge and become increasingly differentiated, ultimately resulting in a population composed exclusively of unisexual males and females (Figure 2C, Geritz et al., 1998; De Jong and Geritz, 2001; Lesaffre et al., 2024, 2025). Initially, the two alleles diverge in both sex allocation components (*z*_0_ and *b*), such that individuals carrying the allele specialising into female function are characterised by higher a baseline (higher *z*_0_) and feminisation (positive *b*), while those carrying the male-specialising allele show a lower baseline and masculinisation (lower *z*_0_ and negative *b*). Thus, as the transition unfolds, individuals carrying the female and male allele exhibit increased allocation to their preferred function with size. As alleles keep diverging, however, they evolve such strongly diverged baselines that the slope components eventually no longer contribute to the expressed phenotype. The slope components are then selected towards zero owing to the cost of plasticity (eq. 2).

### 3.2 Limited seed dispersal counteracts the effects of selfing and can lead to gynodioecy

We next investigated how limited seed dispersal and selfing jointly determine the size-dependent strategies that evolve. As a baseline, we ran simulations under complete outcrossing and no environmental persistence (*α* = 0 and *h* = 0) to isolate the effect of limited seed dispersal (0 *< m <* 1). In line with previous results (Sakai and Sakai, 2003), we found that limited seed dispersal generates a budget effect favouring masculinisation (*b <* 0, Supp. Fig. S2). This is because limited seed dispersal leads a plant’s own offspring to compete with one another for access to resources, which generates diminishing fitness returns to seed production as a result of kin competition (local resource competition; Clark, 1978). This kin competition effect is stronger in larger, more fecund plants, which favours a shift towards male function in these individuals.

We conducted a parameter sweep to investigate how the budgets effects generated by selfing and limited seed dispersal interact to determine the pattern of size-dependent sex allocation that evolves. When inbreeding depression is low (*δ <* 1*/*2), selfing and limited seed dispersal have opposing effects, and which of the two predominates depends on the selfing and dispersal rates: when selfing is rare and seeds mostly remain on their natal patch (low *α* and low *m*), the effect of limited seed dispersal predominates and selection favours masculinisation. Conversely, at high rates of selfing and seed dispersal (*α* and *m* both high), selection is mostly driven by the budget effect of selfing and favours feminisation (Figure 3A). Inbreeding depression weakens the effect of selfing relative to limited seed dispersal, which it does not affect. As a result, increasing inbreeding depression increases the parameter range over which masculinisation evolves (compare panels A and B in Fig. 3).

**Figure 3:**
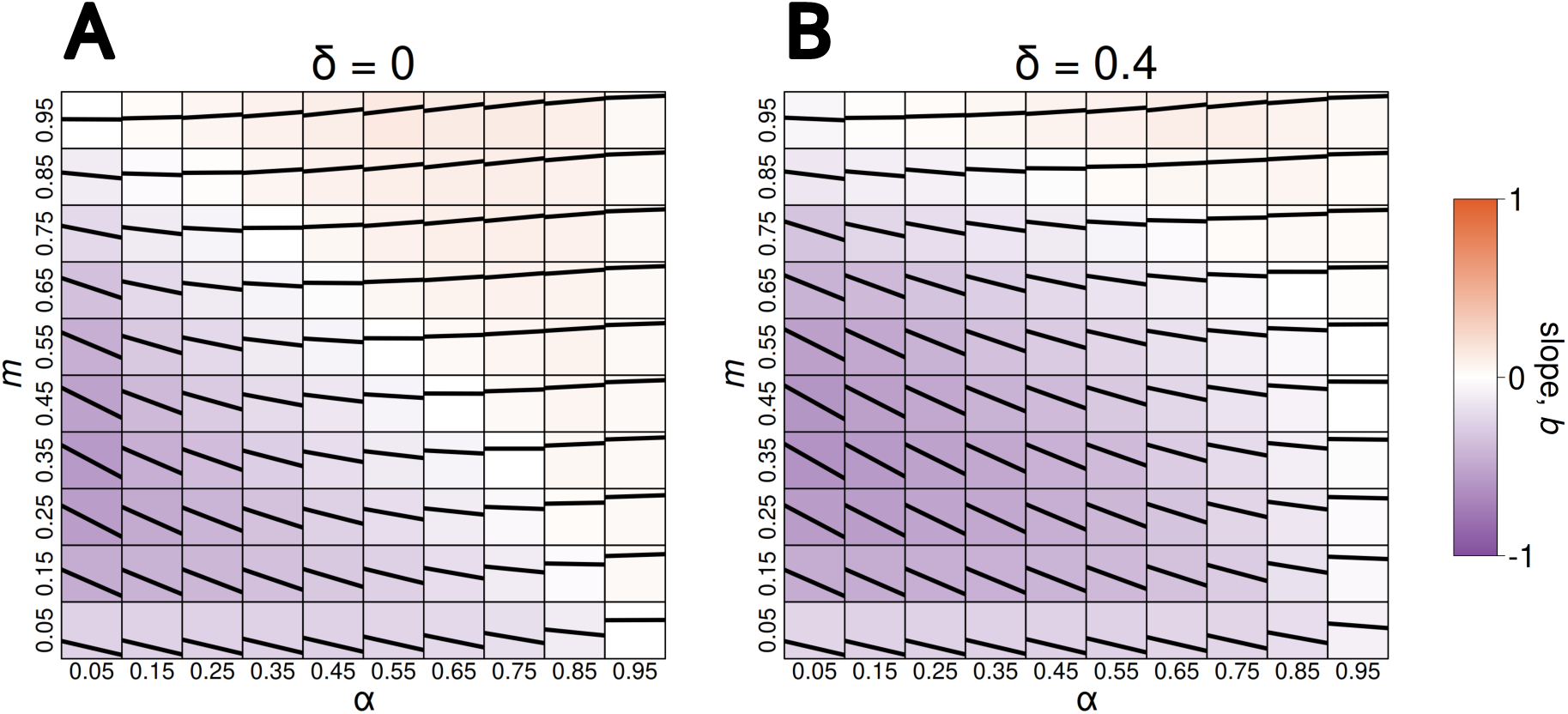
Evolved size-dependent sex allocations as a function of the selfing rate *α* (x-axis) and the seed dispersal rate *m* (y-axis), in the absence of inbreeding depression (*δ* = 0, Panel A) and under moderate inbreeding depression (*δ* = 0.4, Panel B). For each parameter combination, the solid black line shows the evolved reaction norm averaged over the last 5 × 10^5^ generations across three replicates of simulations run for 2 × 10^6^ generations each. The background is coloured according to the slope of the evolved reaction norm, with shades of orange and purple indicating feminisation and masculinisation, respectively. Darker shades indicate stronger size-dependence. Parameters used in all simulations: *N* = 5, 000, *µ* = 0.005, *σ_µ_* = 0.01, *h* = 0, *c* = 0.01.

When inbreeding depression is high (*δ >* 1*/*2), selfing becomes net-detrimental, leading to selection for inbreeding avoidance. In contrast with the complete seed dispersal case (*m* = 1), however, this does not necessarily lead to the emergence of a sexual polymorphism. This is because the diminishing fitness returns to ovule production generated by limited seed dispersal oppose the increasing fitness returns created by inbreeding avoidance, which can stabilise hermaphroditism (Lesaffre et al., 2025). In this case, maleness increases strongly with size because both budget effects select in the same direction under high inbreeding depression (Fig. 4A,B): larger plants tend to self-fertilise more, which reduces their expected fitness returns through female function and so selects for masculinisation.

**Figure 4:**
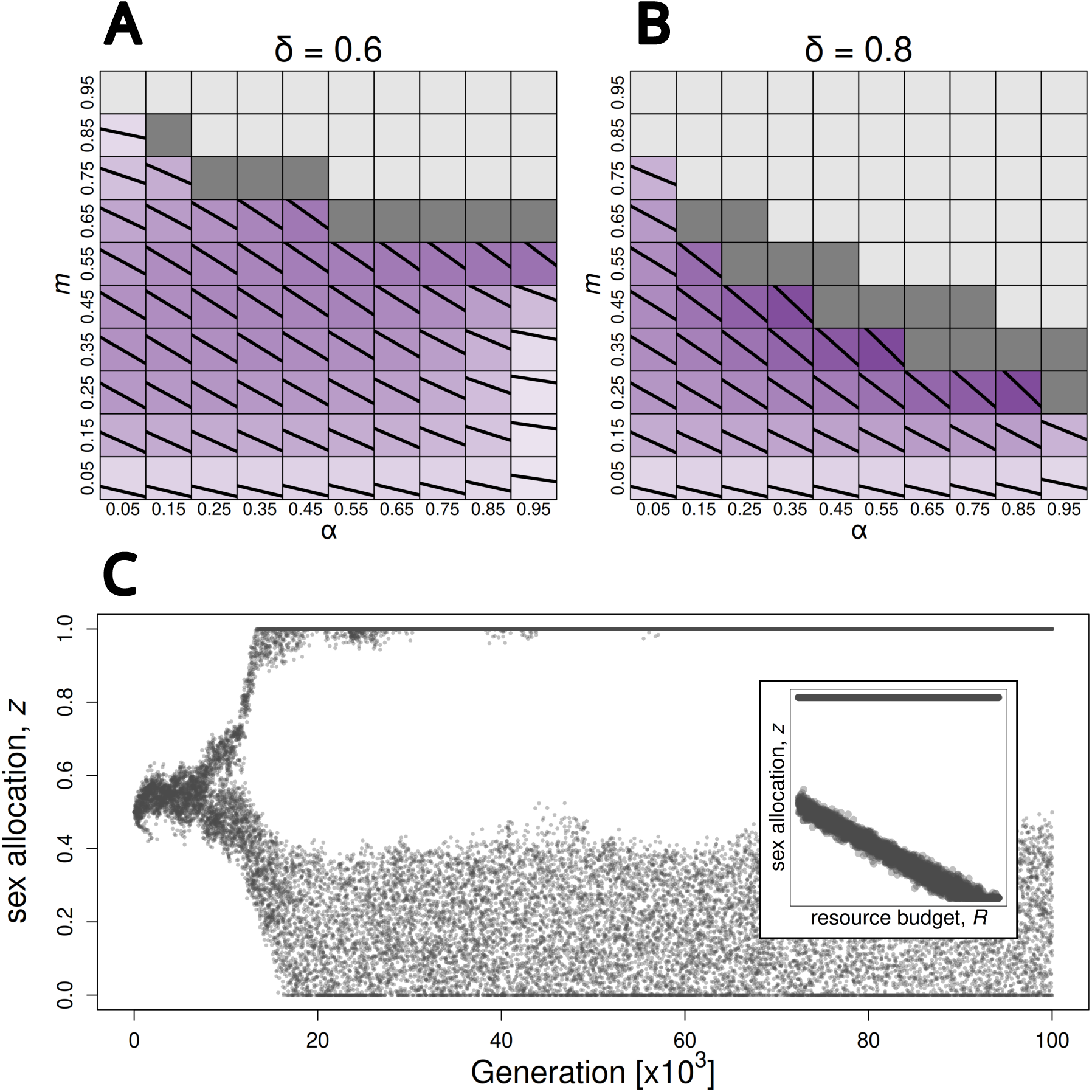
Evolved size-dependent sex allocations under high inbreeding depression. (A) *δ* = 0.6. (B) *δ* = 0.8. As in Fig. 3 above, the solid black line shows the evolved reaction norm, and the background is coloured according to the evolved slope component (the colour scale is the same as Fig. 3). Full light and dark grey squares indicate parameter sets for which dioecy and gynodioecy evolved, respectively. (C) Distribution of phenotypes through time in a simulation where gynodioecy evolved. Each point is an individual randomly sampled from the population. The subplot on the right-hand side shows the phenotype expressed by individuals as a function of their resource budget from generation 80 × 10^3^ to 100 × 10^3^. Parameters used in all simulations: *µ* = 0.005, *σ_µ_* = 0.01, *m* = 1, *h* = 0, *c* = 0.01. Population size was set to *N* = 5 × 10^3^ as a baseline, but was increased to *N* = 10^5^ in cases where simulations with *N* = 5 × 10^3^ yielded ambiguous results (e.g., repeated but aborted branching events).

Sexual polymorphism does nevertheless emerge above a threshold seed dispersal rate that depends on the selfing rate *α* and inbreeding depression *δ* (Fig. 4A,B). When the dispersal rate is sufficiently high relative to this threshold, fully separate sexes evolve, as before. For dispersal rates above but close to the threshold, on the other hand, gradual evolution leads to gynodioecy, where unisexual females coexist with male-biased hermaphrodites that display masculinisation (Fig. 4C). This is likely because *(i)* limited seed dispersal reduces the fitness returns gained by the male-biased morph from fertilising ovules on females, as their non-dispersed seeds experience intense competition for recruitment and are therefore more likely to fail to settle; and *(ii)* relatively fewer seeds reach the habitat patches occupied by malebiased individuals under limited dispersal, so that it is worthwhile for them to invest some resources in producing seeds, as those have better chances of settling on their natal patch.

### 3.3 Environmental persistence favours feminisation under seed limited dispersal

So far, we have assumed that patch qualities are randomly assigned each generation, in keeping with most theory on the evolution of size-dependent sex allocation (e.g., De Jong et al., 1999; Zhang and Jiang, 2002; Sakai and Sakai, 2003; Sato, 2004; Cadet et al., 2004). Yet, patch-quality differences may often persist across several generations. For example, some patches of habitat may consistently be better exposed to sunlight, give easier access to underground water or be more nutrient-rich. To investigate the influence that such environmental persistence might have on sex allocation evolution, we ran simulations where we varied the probability *h* that a patch remains of the same quality in the next generation.

Under complete outcrossing (*α* = 0), our simulations reveal that environmental persistence (*h >* 0) favours feminisation when seed dispersal is limited (Fig. 5A). This is because it generates an additional budget effect: seeds produced by larger plants germinate on higher-quality patches on average, because non-dispersed seeds are more likely to experience the same growing environment as their mother (Guillon et al., 2006). This increases their reproductive value, defined as their relative contribution to the population in the distant future (Taylor, 1990; Lehmann and Rousset, 2014), and so promotes greater allocation to female function in large plants. Conversely, seeds produced by smaller plants have a lower reproductive value than average, so that it is more worthwhile for small individuals to allocate more to pollen production and fertilise ovules on other, potentially better provisioned plants.

**Figure 5:**
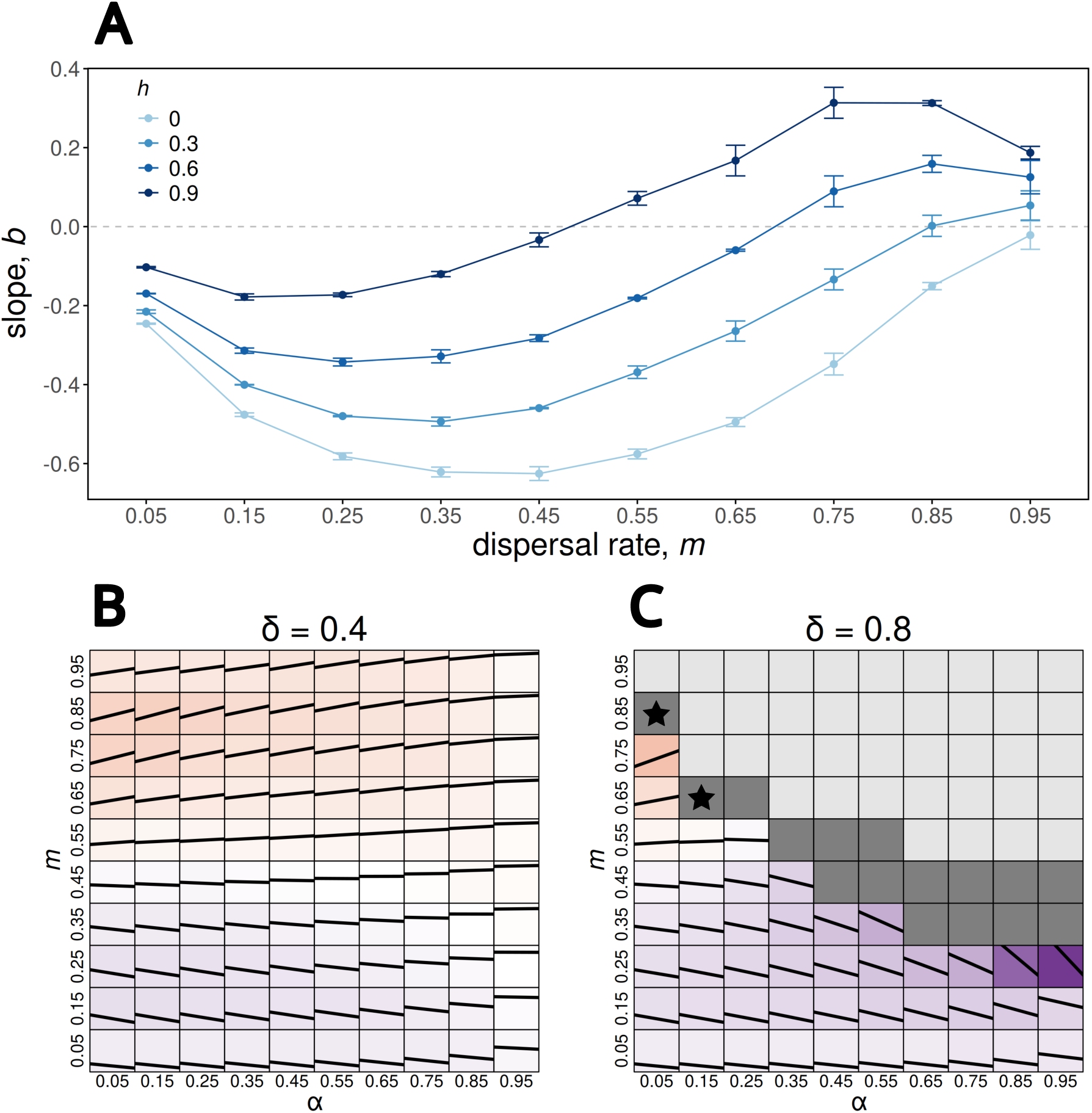
Effect of environmental persistence on the evolution of size-dependent sex allocation. (A) Evolved slope components as a function of the seed dispersal rate (*m*) under complete outcrossing (*α* = 0) for different levels of environmental persistence (*h*). Slopes were measured over the last 5 × 10^5^ generations of simulations run 2 × 10^6^ generations, and averaged across replicates. (B) Evolved sex allocations under moderate inbreeding depression (*δ* = 0.4) and high environmental persistence (*h* = 0.9), as a function of the selfing and dispersal rates. As above, the solid black lines show the evolved reaction norms, and the background is coloured according to the direction of size-dependence (same scale as previous figures). (C) Evolved sex allocations under high inbreeding depression (*δ* = 0.8) and environmental persistence (*h* = 0.9). Grey tiles indicate parameter combinations for which either dioecy (light grey) or gynodioecy (dark grey) evolved. Black stars mark parameter combinations for which environmental persistence qualitatively alters the sexual polymorphism that evolves (see main text). Parameters used in all simulations: *µ* = 0.005, *σ_µ_* = 0.01, *c* = 0.01. Population size was set to *N* = 5 × 10^3^ and increased to *N* = 10^5^ when necessary.

Limited seed dispersal thus has two opposing effects on size-dependent sex allocation under environmental persistence. On the one hand, decreasing the seed dispersal rate increases the likelihood that a plant’s own seeds will inherit its own patch, which selects for feminisation. On the other hand, it increases the intensity of local resource competition, and so selects for masculinisation. The kin competition effect tends to dominate when environmental persistence and the seed dispersal rate are low (low *h* and *m*, Fig. 5A), leading larger individuals to become more male. Conversely, the effect of environmental persistence dominates and causes limited seed dispersal to favour feminisation when environmental persistence and seed dispersal are high (high *h* and *m*).

We also investigated how the interaction between the budget effects of selfing and limited seed dispersal is modified by environmental persistence under weak and strong inbreeding depression. Under low inbreeding depression (*δ <* 1*/*2), environmental persistence substantially extends the range of parameters for which feminisation evolves (compare Fig. 2B with Fig. 5B). Under high inbreeding depression, the population either remains monomorphic or evolves a sexual polymorphism owing to selection for inbreeding avoidance, as before. When the population remains monomorphic, environmental persistence promotes feminisation, similar to the low inbreeding depression case, resulting in weaker size-dependence overall as selfing and local resource competition are both favouring the opposite pattern of size-dependence here (compare Fig. 4B with Fig. 5C).

When sexual polymorphism emerges, meanwhile, we find that environmental persistence alters it in two main ways. First, in conditions leading to gynodioecy, we find that environmental persistence causes the hermaphroditic morph to evolve feminisation instead of masculinisation (Supp. Fig. S3A). This evolves for the same reason as in the monomorphic case: when the environment persists across generations, well-provisioned hermaphrodites are selected to allocate more to their female function because their own progeny is then more likely to inherit their high-quality patch. Second, for some combinations of high seed dispersal rates and low selfing rates (marked with a star in Fig. 5C), we find that the female morph no longer remains strictly unisexual. Instead, we find that while it remains heavily female-biased, it now allocates some resources to pollen production when growing in poor conditions (Supp. Fig S3B). This is likely because environmental persistence reduces expected fitness returns of seeds relative to pollen for individuals growing on low-quality patches.

## 4 Discussion

De Jong et al. (1999) were the first to investigate the influence of selfing on the evolution of size-dependent sex allocation. Using a model where the selfing rate increases with the individual’s pollen production, but is independent of the pollen production of other individuals in the population, they found that selfing favours feminisation. They suggested that this pattern could be explained by the greater difficulty for larger plants to avoid geitonogamous pollen transfers. Here, we showed that competing selfing, where the selfing rate depends on other individuals’ pollen production as well, favours a similar relationship between size and sex allocation. However, we propose a different explanation for this pattern. Rather than reflecting selection to avoid accidental selfing, we argue that, when inbreeding depression is sufficiently weak for selfed seeds to yield greater average fitness returns than outcrossed seeds (*δ <* 1*/*2 in our model), selection favours larger plants that allocate more to female function because doing so allows them to produce more self-fertilised seeds overall. When inbreeding depression is strong (*δ >* 1*/*2), on the other hand, selection favours either the emergence of separate sexes (De Jong and Geritz, 2001), or masculinisation when other ecological mechanisms select against sexual specialisation.

Two mechanisms that are known to select against sexual specialisation are the limited dispersal of pollen and seed, which lead related individuals to compete with one another for access to local mates and resources, respectively, generating diminishing fitness returns to specialising as male and female (Fromhage and Kokko, 2010; Lesaffre et al., 2025). In addition to stabilising hermaphroditism, limited seed and pollen dispersal have been shown to influence the evolution of size-dependent sex allocation (Sakai and Sakai, 2003). Limited pollen dispersal favours feminisation, meanwhile limited seed dispersal favours masculinisation. We investigated how selfing and limited seed dispersal interact to determine the size-dependent sex allocation strategy that evolves, as these two mechanisms favour opposite patterns of size-dependence. We showed that which of the two predominates depends on the selfing and seed dispersal rates. In the absence of inbreeding depression, the effect of selfing predominates and feminisation evolves when the selfing rate exceeds the proportion of seeds that remain on their natal patch (1 − *m < α*). Conversely, masculinisation evolves whenever the seed dispersal rate falls below this threshold. As inbreeding depression increases, however, masculinisation evolves over a wider region of the parameter space because inbreeding depression reduces the fitness benefits of selfing, and so weakens its influence on size-dependent sex allocation. This result suggests that plants should more often display masculinisation in the presence of inbreeding depression, unless they exhibit high selfing and seed dispersal rates. However, available empirical evidence supports the opposite pattern: feminisation is in fact far more frequent (e.g., De Jong and Klinkhamer, 1989; Klinkhamer et al., 1997; De Jong and Klinkhamer, 2005).

Explaining the apparent prevalence of feminisation in flowering plants therefore requires considering additional mechanisms. In particular, the prediction that masculinisation may evolve more readily in the presence of inbreeding depression rests on two assumptions that are unlikely to hold in nature. First, it relies on the assumption that patch quality changes randomly each generation. However, fine-scale differences in the environmental conditions experienced by different individuals in a plant population might persist over several generations. Such persistence could arise, for instance, from geological features that generate spatial variation in sunlight exposure, water availability or subsoil composition, as well as from disturbances that occur more frequently in some patches than in others. We relaxed this assumption in the last part of our analysis, and showed that the persistence of environmental heterogeneity generates a budget effect that favours increased allocation to female function in larger plants when seed dispersal is limited, because higher seed production in these plants then increases the probability that their high-quality patch will be passed on to their own offspring (Guillon et al., 2006). This feminising effect dominates over the masculinising effect of local resource competition at higher dispersal rates. Thus, persistent differences in the micro-environments experienced by individuals might help explain the prevalence of feminisation in plants. Second, we assumed that exported pollen is distributed equally among other individuals in the population according to a classical mass-action model (Holsinger, 1991), so that pollen dispersal is not limited. Yet, empirical evidence points to pollen transfers often occurring over short distances (Webb, 1998; Vekemans and Hardy, 2004; Torices et al., 2026), suggesting that limited pollen dispersal may be a pervasive force selecting for feminisation in plants. In the future, this assumption could be relaxed by extending our model to include an explicit spatial structure (Fromhage and Kokko, 2010).

We assumed that resource budget differences between plants were purely environmentally determined. In reality, however, the amount of resources acquired by a plant may also depend on its genotype and in particular on its load of deleterious mutations: individuals carrying more mutations might exploit available resources at a slower rate, and so perform less well than less-loaded individuals even when they develop on a better patch. Explicit modelling of such a mutation load would mechanically generate inbreeding depression, which we treated as a parameter in this study. This would allow one to investigate the effect of purging on the evolution of size-dependent sex allocation, which has not been considered so far despite it being recognized as an important mechanism in mating-system evolution (Lande and Schemske, 1985; Charlesworth et al., 1990, 1991; Abu Awad and Roze, 2020). In addition, resource budget would then become an indicator of genetic quality, which may generate new selective pressures on size-dependent sex allocation. Selection broadly favours the evolution of condition-dependent self-fertilisation (Lesaffre et al., 2026b), whereby individuals adjust their selfing rate to their biological condition (defined as an individual’s overall vitality owing to its genetic background and environment; Wilson and Nussey, 2010; Forsythe et al., 2021). In short, this plasticity for the selfing rate is favoured as an escape mechanism from more deleterious backgrounds: from the point-of-view of a modifier allele found in a heavily loaded genetic background, outcrossing is a better mating strategy than selfing because it gives the allele a chance to recombine into a better background (Lesaffre et al., 2026b). If sex allocation influences the selfing rate, as found in a number of species (Harder and Barrett, 1995; Vrieling et al., 1999; Chen and Pannell, 2024, 2025b), then selection for condition-dependent self-fertilisation could affect the evolution of size-dependent sex allocation in ways to remain to be explored.

## Acknowledgements

The authors thank the Swiss National Science Foundation for funding (SNF grants 310030 220139 to CM and PZ00-3 232857 to TL).

## Statement of Authorship

SM and TL conceived the study. SM prepared the simulation program and carried out the analysis with input from TL. SM and TL wrote the first draft. All authors contributed to revisions and approved the final manuscript. TL and CM acquired the funding.

## Data and Code Availability

Simulation data and code are publicly available at 10.5281/zenodo. 24459591.

