## Supplementary Material for "The effects of selfing, limited seed dispersal and environmental persistence on the evolution of size-dependent sex allocation"

#### Table of contents

**A Individual-based simulation program** **2**

**B Supplementary figures** **5**

#### Appendix A

### Individual-based simulation program

In this appendix, we describe the individual-based simulation program that we used to obtain the results presented in the paper. The program is coded in C++20, and is publicly available along with the simulation data at: `10.5281/zenodo.24459591`.

The program simulates a population of $N$ haploid individuals, each characterised by their genetically encoded value for two traits: the baseline ($z_0$) and slope ($b$) components of sex allocation. The traits are assumed to be encoded by two fully linked loci following a continuum-of-alleles model. Together, they determine the proportion of resources that an individual allocates to its female and male function as a function of its resource budget $R$. The population evolves in an environment composed of $N$ patches hosting one adult plant each. Each patch is characterised by its quality, which determines the resource budget $R$ available to the individual growing on it.

**Initialisation.** At the beginning of a simulation, the population is initialised with $N$ individuals that all share the same baseline and slope sex allocation components $z_{0,\text{init}}$ and $b_{\text{init}}$. Patch qualities are initialised by sampling $N$ values from a uniform distribution $\mathcal{U}[0, 1]$.

The simulation then runs for $t_{\text{max}}$ generations, and each generation proceeds as follows.

**Sex allocation, fecundities, and effective selfing and local recruitment probabilities.** We first determine the sex allocation expressed by each individual as a function of its trait values and resource budget. For individual $i$, this is

$$z_i = \begin{cases} 0 & \text{if } z_{0,i} + b_i R_i < 0, \\ z_{0,i} + b_i R_i & \text{if } 0 \leqslant z_{0,i} + b_i R_i \leqslant 1, \\ 1 & \text{if } z_{0,i} + b_i R_i > 1. \end{cases} \tag{A1}$$

From these, we determine individual pollen productions as

$$M_i = R_i(1 - z_i)\exp\left(-c{b_i}^2\right) \tag{A2}$$

for individual $i$, and compute the total pollen production in the population as $M_{\text{tot}} = \sum_{i=1}^{N} M_i$.

Using pollen productions, we then compute the selfing rate of each individual in the population as

$$s_i = \frac{\alpha M_i}{\alpha M_i + \dfrac{1-\alpha}{N-1}(M_{\text{tot}} - M_i)} \tag{A3}$$

for individual $i$. The selfing rate amongst the surviving seeds of individual $i$, which accounts for seed mortality owing to inbreeding depression, can then be computed as

$$s_i^{\delta} = \frac{s_i(1-\delta)}{1 - s_i\delta}. \tag{A4}$$

Meanwhile, the female fecundity of individual $i$ is

$$F_i = R_i z_i \exp\left(-c{b_i}^2\right). \tag{A5}$$

The number of viable seeds produced by individual $i$ after accounting for inbreeding depression is then obtained as

$$F_i^{\delta} = F_i(1 - s_i\delta), \tag{A6}$$

so that the probability that a patch will be occupied by an offspring of the local individual in the next generation is

$$P_i = \frac{F_i^{\delta}(1-m)}{F_i^{\delta}(1-m) + \dfrac{m}{N-1}\left(F_{\text{tot}}^{\delta} - F_i^{\delta}\right)}, \tag{A7}$$

where $F_{\text{tot}}^{\delta} = \sum_{i=1}^{N} F_i^{\delta}$ is the total number of viable seeds produced by individuals in the population.

**Offspring production.** For each patch $i$, we first determine whether the recruited seed will be a migrant or a local using probabilities given in eq. (A7). If it is a migrant, we then sample a mother amongst the

$N-1$ individuals located on other patches according to their effective female fecundities: each individual $j \neq i$ has a probability $F_j^\delta / \sum_{k \neq i} F_k^\delta$ of being sampled. If it is a local, then the mother is individual $j = i$. Once the mother has been chosen, the recruited seed is either self-fertilised or outcrossed with a probability given by the mother's effective selfing rate $s_j^\delta$. If it is a selfed, the seed inherits the mother's haplotype. If it is outcrossed, it inherits the mother's haplotype with probability $1/2$. Otherwise, we sample a father according to male fecundities (eq. A2), so that the probability that individual $k \neq j$ is chosen is $M_k / \sum_{\ell \neq j} M_\ell$, and the seed inherits its haplotype.

**Mutation and environmental change.** Each recruited seed then undergoes mutation with probability $\mu$, in which case independent random deviates are added to its baseline and slope components $z_0$ and $b$. Mutation effects are sampled in a normal distribution $\mathcal{N}(0, \sigma_\mu)$. The patch occupied by each seed either remains of the same quality with probability $h$, or a new resource budget $R$ is sampled in a uniform distribution $\mathcal{U}[0, 1]$.

**Measurements.** Every $t_{\text{mes}}$ generations, we record the genetic values of baseline and slope components, the resource budget and the expressed sex allocation of $n_{\text{mes}}$ individuals in separate output files.

#### Appendix B

### Supplementary figures

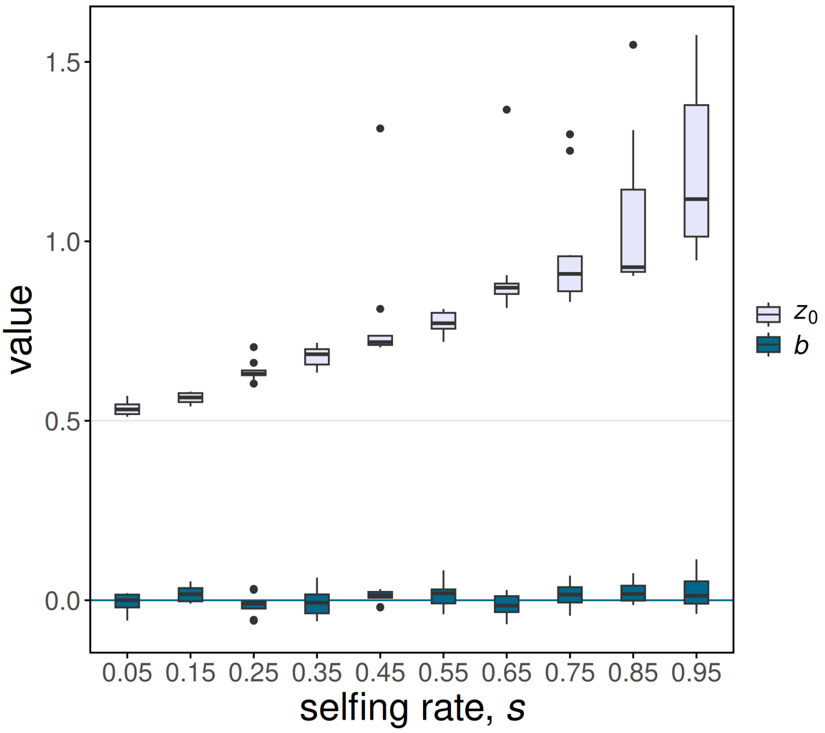


**Figure S1:** Effect of a fixed selfing rate $s$ on the evolution of size-dependent sex allocation. Boxplots represent the distribution of the average baseline and slope components of sex allocation, $z_0$ and $b$, in the last $5 \times 10^5$ generations of ten replicate simulations run for a total of $10^6$ generations each. Horizontal lines mark the values of $z_0$ and $b$ at the start of the simulations. With a fixed selfing rate, the slope component $b$ remains close to zero, whereas the baseline $z_0$ evolves towards female-biased values, as expected. Parameters used in all simulations: $N = 5 \times 10^3$, $\mu = 0.005$, $\sigma_\mu = 0.01$, $m = 1$, $h = 0$, $\delta = 0$, $c = 0.01$.

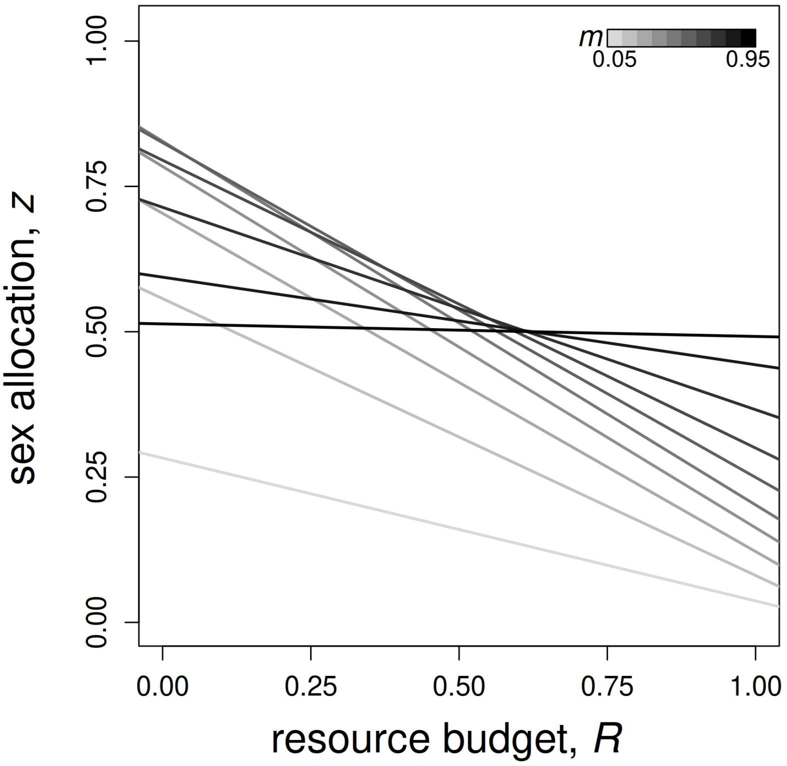


**Figure S2:** Effect of limited seed dispersal on the evolution of size-dependent sex allocation under complete outcrossing and in the absence of environmental persistence ($\alpha = h = 0$). Each line represents the evolved size-dependent sex allocation, obtained by averaging across replicates over the last $5 \times 10^5$ generations of simulations, as in other plots. Darker shades of grey indicate higher dispersal rates.

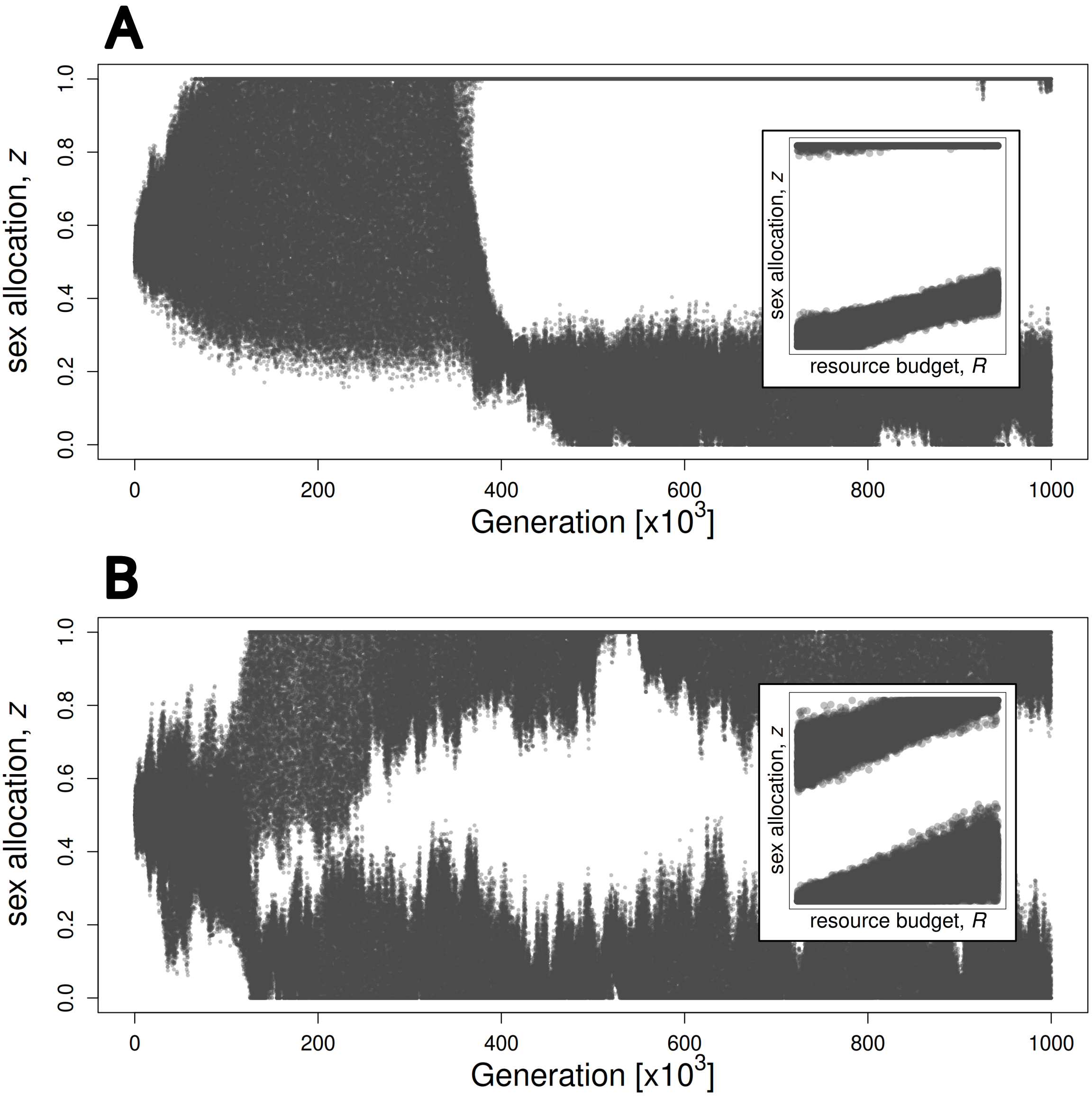


**Figure S3:** Effect of environmental persistence on the sexual polymorphism that evolves under high inbreeding depression. In both panels, each point corresponds to a randomly sampled individual in the population. The main plot shows how the phenotypic distribution of the population changes through time, and the subplot shows individual sex allocations as a function of resource budget from generation $8 \times 10^5$ to $10^6$. (A) Gynodioecy where the hermaphroditic morph shows feminisation. Parameters used in this simulation: $\alpha = 0.55$, $m = 0.45$, $\delta = 0.8$ and $h = 0.9$. (B) Evolution of a form of subdioecy, where the female-biased morph allocates more to pollen production when growing in poor conditions. Parameters used in this simulation: $\alpha = 0.05$, $m = 0.85$, $\delta = 0.8$ and $h = 0.9$. Other parameters: $N = 10^5$, $\mu = 0.005$, $\sigma_\mu = 0.01$, $c = 0.01$.
